# An interferon-like off-switch in invertebrates: Vago4 downregulates Domeless to restrain JAK/STAT signaling

**DOI:** 10.64898/2026.10.26.761873

**Authors:** Linwei Yang, Jianhui He, Alian Liao, Deyu Han, Wanli Lu, Hongliang Zuo, Jianguo He, Xiaopeng Xu

## Abstract

Whether invertebrates possess a functional interferon (IFN)-like system remains controversial, as canonical IFN ligands and their receptors are absent from invertebrate genomes. Although Vago proteins in *Drosophila* and shrimp are compelling functional analogs of IFN, direct evidence for a canonical cytokine–receptor axis and, importantly, for how such signaling is restrained to prevent excessive immune activation, has been lacking. Here, we identify Vago4 as a bona fide ligand of Domeless, the canonical type I cytokine receptor upstream of JAK/STAT signaling. Vago4 binds the extracellular CBM1 and CBM2 domains of Domeless, triggering receptor endocytosis followed by lysosomal trafficking and K27-linked ubiquitination-dependent proteasomal degradation. This ligand-induced receptor downregulation serves as a negative feedback mechanism that restrains excessive JAK/STAT activation. During acute *Vibrio* infection, Vago4 silencing accelerated shrimp mortality, whereas recombinant Vago4 alleviated inflammatory lesions and oxidative stress, enhanced phagocytosis, and improved survival, phenocopying JAK or STAT knockdown. Thus, by downregulating Domeless, Vago4 limits immunopathology and defines an IRF–Vago4–Domeless–JAK/STAT IFN-like axis in invertebrates, offering new insight into the evolution of IFN-mediated immune regulation.

**Author Summary:** In vertebrates, interferons are important immune-regulating cytokines that play key roles in antiviral and antibacterial immunity, but their excessive activation can also damage the body. Whether invertebrates possess an interferon-like immune system has remained an open question, given that they lack the classic interferon genes and receptors present in vertebrates. A protein called Vago4, identified in arthropods, was found to function like interferon. We showed that Vago4 binds to Domeless, a cell-surface receptor that serves as the invertebrate counterpart of the cytokine receptors through which interferons signal in vertebrates. This binding causes Domeless to be taken into the cell and degraded, thereby limiting immune signaling and preventing the immune response from becoming too strong. During acute infection with *Vibrio* bacteria, shrimp in which Vago4 was suppressed died faster. However, after injection of the Vago4 protein, shrimp showed reduced inflammation, lower harmful oxidative stress, enhanced bacterial engulfment, and improved survival. Our findings show that invertebrates have an interferon-like “off-switch” that helps maintain immune balance.

## 1. Introduction

The JAK/STAT pathway, composed of cytokine receptors, Janus kinases (JAKs), and signal transducers and activators of transcription (STATs), is a central mediator of cytokine and growth factor signaling and plays critical roles in regulating cell proliferation, differentiation, apoptosis, and immune responses(Murray, 2007; Stark and Darnell, 2012). Ligand-mediated dimerization of cell-surface receptors, such as interferon (IFN) receptor (IFNAR) and epidermal growth factor receptor (EGFR), brings receptor-associated JAKs into close proximity and enables their activation via reciprocal tyrosine phosphorylation. Activated JAKs subsequently phosphorylate specific tyrosine residues on the receptor, creating docking sites for STAT proteins. STATs are recruited through their SH2 domains and phosphorylated by JAKs. Activated STATs form homo- or heterodimers, translocate to the nucleus, and bind to specific DNA elements to regulate target gene transcription(Murray, 2007). In invertebrates, Domeless is the only identified membrane receptor homologous to vertebrate type I cytokine receptors(Brown *et al*., 2001). It contains two cytokine-binding modules and three fibronectin type III domains and shares high structural similarity with the vertebrate interleukin-6 receptor (IL-6R)(Yan *et al*., 2015). In *Drosophila*, Domeless directly binds three Unpaired (UPD1–3) ligands, activating the JAK kinase Hopscotch and the STAT protein STAT92E, thereby regulating embryonic development and tracheal morphogenesis(Brown *et al*., 2001). In crustaceans, Domeless likewise activates the JAK/STAT pathway and participates in antibacterial and antiviral immune responses (Sun *et al*., 2017; Yan *et al*., 2015).

In vertebrates, IFNs are key upstream cytokines in the JAK/STAT pathway and play central roles in antibacterial and antiviral immunity(Mertowska *et al*., 2023; Raftery and Stevenson, 2017). Due to the absence of sequences homologous to IFNs and their receptors in invertebrate genomes, IFN-mediated signaling pathways were long thought to be lacking in invertebrates(Arbouzova and Zeidler, 2006; Obbard and Dudas, 2014). This view has now been challenged by the identification of Vago, an invertebrate cytokine with IFN-like functions(Li *et al*., 2015; Paradkar *et al*., 2012). Vago is a secreted protein containing a von Willebrand factor C (VWC) domain and is widely found in arthropods, including fruit flies, mosquitoes, and shrimp(Deddouche *et al*., 2008; Li *et al*., 2015; Paradkar *et al*., 2012). Although Vago lacks sequence homology with IFNs, it can induce an antiviral state by activating the JAK/STAT pathway and is therefore considered a functional analog of IFNs(Paradkar *et al*., 2012). In *Culex* mosquitoes, Vago suppresses West Nile virus (WNV) infection via activation of the JAK/STAT pathway(Paradkar *et al*., 2012). In shrimp, the DNA-sensing STING-IFN regulatory factor (IRF) axis activates Vago4 expression, which in turn triggers JAK/STAT signaling, forming a regulatory cascade analogous to the mammalian STING-IRF-IFN-JAK/STAT axis and further supporting that Vago functions as an interferon analog in invertebrates(Gao *et al*., 2019; Li *et al*., 2015). In Pacific white shrimp *Penaeus vannamei*, Vago is also implicated in antibacterial response by regulating the expression of antimicrobial peptides(Boonchuen *et al*., 2022). Moreover, Vago has been reported to function as a pattern recognition receptor (PRR), directly recognizing pathogens and activating the JAK/STAT pathway(Qin *et al*., 2023; Qin *et al*., 2020; Qin *et al*., 2022). Notably, in *Marsupenaeus japonicus*, Vago-L activates the JAK/STAT pathway through a non-classical receptor, integrin, rather than the canonical receptor Domeless(Gao *et al*., 2021). However, the functional link between Vago and the classic JAK/STAT receptor Domeless remains unclear.

The activity of the JAK/STAT pathway is tightly controlled. In mammals, excessive JAK/STAT activation is a major driver of inflammation, sepsis, and cytokine storms(Li *et al*., 2026; Nie *et al*., 2025). Recent studies in an acute *Vibrio* infection model of *P. vannamei* have also shown that excessive activation of the JAK/STAT pathway, rather than the NF-κB signaling cascades, is a primary cause of inflammation-like lesions in invertebrates(Zuo *et al*., 2025). Negative regulatory mechanisms of the JAK/STAT pathway mediated by suppressor of cytokine signaling (SOCS), protein inhibitor of activated STAT (PIAS), and dual-specificity phosphatase 14 (DUSP14) are essential for maintaining immune homeostasis(Luo *et al*., 2024; Niu *et al*., 2018; Wang *et al*., 2016). Furthermore, in IFN-mediated activation of the JAK/STAT pathway in vertebrates, IFNAR can be internalized and degraded in lysosomes, thereby terminating signal transduction and preventing excessive inflammatory responses(Chmiest *et al*., 2016). During inflammation or cytokine storms, IFNs can promote the endocytosis, ubiquitination, and lysosomal degradation of IFNAR1, limiting the sustained activation of STAT1 and STAT2(Chmiest *et al*., 2016). For example, IFNα induces phosphorylation of IFNAR1 at Ser535, which facilitates recruitment of the E3 ubiquitin ligase β-Trcp2, leading to ubiquitination and proteasomal degradation of IFNAR1(Bhattacharya *et al*., 2010). In *Drosophila*, the receptor Domeless has also been shown to undergo endocytosis and lysosomal degradation, thereby attenuating JAK/STAT signaling activity(Hou *et al*., 2015; Stec *et al*., 2013). However, as a functional IFN analog in invertebrates, the role of Vago in excessive JAK/STAT activation and the consequent immune pathology remains unclear.

The current study revealed a Vago-mediated negative regulatory mechanism of JAK/STAT signaling during immune overactivation. We demonstrated that Vago4 functions as an inducible ligand that binds the Domeless receptor, promoting its internalization and subsequent lysosomal degradation. This process restores immune homeostasis by attenuating excessive JAK/STAT signaling, preventing host death caused by acute *Vibrio* infection. These findings elucidate a ligand-mediated mechanism of receptor internalization and degradation in invertebrate JAK/STAT signaling and, more importantly, identify Vago as an unconventional negative regulator in immune overactivation, paralleling mammalian IFNs, offering new insights into the maintenance of immune homeostasis in invertebrates.

## 2. Results

### 2.1. Vago4 inhibits the JAK/STAT signaling pathway

STAT nuclear translocation is a critical indicator of JAK/STAT signaling pathway activation. Immunofluorescence analysis revealed that both poly(I:C) and *V. parahaemolyticus* stimulation significantly induced STAT nuclear translocation (Fig. S1). In addition, varying doses of *V. parahaemolyticus* infection produced significant differences in STAT nuclear translocation dynamics. In shrimp challenged with a low dose of *V. parahaemolyticus* (NVC, 5 × 10⁵ CFU), STAT nuclear translocation exhibited a transient peak at 1 h post-infection and returned to baseline levels by 4 h. (Fig. 1D). In contrast, high-dose *V. parahaemolyticus* challenge (EVC, 1.5 × 10⁶ CFU) sustained persistent STAT nuclear translocation throughout the 1–4 h period, and STAT activation levels were also higher than in the NVC group (Fig. 1D). These findings confirm that acute *V. parahaemolyticus* infection induces an excessive activation of the JAK/STAT pathway, consistent with previous reports(Zuo *et al*., 2025).

**Figure 1.**
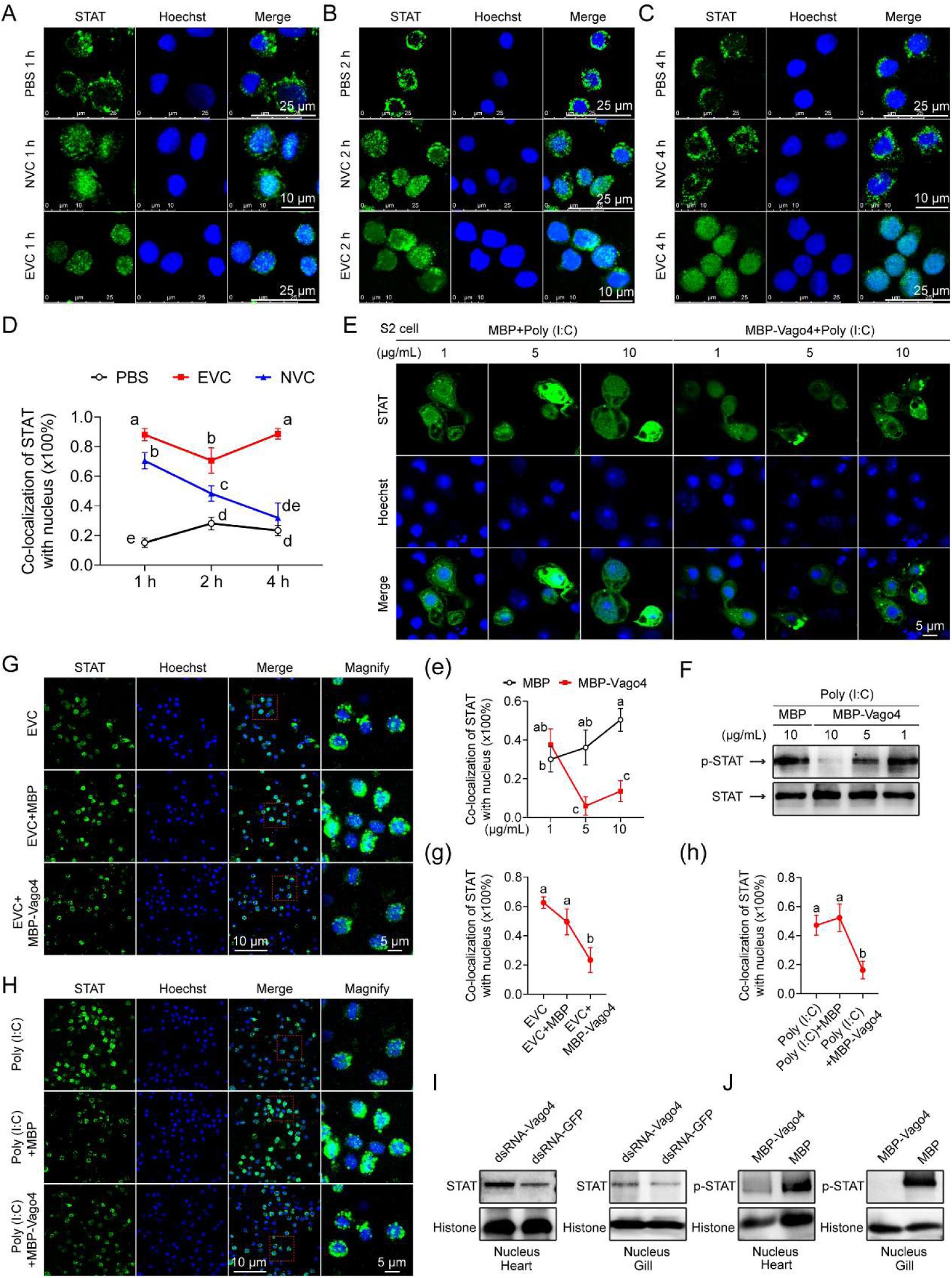
Vago4 suppresses JAK/STAT signaling activation. (A–C) Immunofluorescence analysis of STAT nuclear translocation in shrimp hemocytes at 1 (A), 2 (B), and 4 (C) h post-acute bacterial infection. NVC, the normal *V. parahaemolyticus* challenge, 5 × 10⁵ CFU. EVC, the excessive *V. parahaemolyticus* challenge, 1.5 × 10⁶ CFU. (D) Quantitative analysis of STAT–nucleus colocalization in hemocytes using WCIF ImageJ software. (E–F) Immunofluorescence and Western blot analyses showing the effects of increasing concentrations of recombinant Vago4 (1, 5, or 10 μg/mL) on STAT nuclear translocation and phosphorylation in S2 cells together with 0.5 μg/mL poly(I:C) for 4 h. (G–H) Immunofluorescence analysis of the effects of recombinant Vago4 treatment (10 μg, 4 h) on STAT nuclear localization in shrimp hemocytes together with EVC or poly(I:C) stimulation (5 μg). (I–J) Western blot analysis of nuclear STAT levels in shrimp heart and gill tissues following *Vago4* silencing (10 μg dsRNA/shrimp, 48 h) or recombinant Vago4 treatment (10 μg, 4 h). (e, g, and h) Quantitative analysis of STAT–nucleus colocalization in panels (E), (G) and (H), respectively, using WCIF ImageJ software. Data are shown as mean ± SD. Statistical differences were determined by one-way ANOVA (D, e, g, and h). Values with different letters (D, e, g, and h) indicate significant differences. (*P* < 0.05).

Due to the lack of shrimp cell lines, *Drosophila* S2 cells were for subsequent mechanistic studies. Immunofluorescence analysis confirmed that poly(I:C) significantly induced STAT nuclear translocation in S2 cells (Fig. S2). However, treatment with recombinant Vago4 protein inhibited poly(I:C)-induced STAT nuclear translocation in S2 cells (Fig. 1E and e), and inhibited STAT phosphorylation in a dose-dependent manner (Fig. 1F). This inhibitory effect was further confirmed *in vivo*. Exogenous Vago4 protein treatment significantly reduced EVC- or poly(I:C)-stimulated STAT nuclear translocation in hemocytes (Fig. 1G, g, H and h). Consistently, silencing of *Vago4* gene significantly enhanced STAT nuclear translocation in gill and heart compared to controls (Fig. 1I), while exogenous Vago4 protein treatment showed the opposite effect (Fig. 1J). These results suggest that Vago4 acts as a negative regulator of the JAK/STAT pathway in the context of immune overactivation.

### 2.2. Vago4 interacts with Domeless

The three-dimensional structure of the Vago4–Domeless protein complex was analyzed using AlphaFold 3 and the interaction interface was visualized using PyMOL(Abramson J *et al*., 2024). As shown in Fig. 2A, Vago4 (red) interacted with Domeless (purple) at specific regions. The enlarged view further revealed a hydrogen-bonding network (indicated by dashed lines) formed by multiple key amino acid residues at the interaction interface, providing a structural basis for the stable association between the two proteins. Immunofluorescence analysis revealed significant co-localization between Vago4 (red) and Domeless (green) (Fig. 2B, arrows). Co-IP assays confirmed the interaction between these two proteins (Fig. 2C). The extracellular region of Domeless contains two conserved cytokine-binding homology modules, CBM1 and CBM2, which are structurally similar to those found in the interleukin-6 (IL-6) receptor family (Fig. 2D). Distinct binding interfaces and extensive inter-residue interactions were identified in the docking models of Vago4 with Domeless-CBM1 (pink) and Domeless-CBM2 (blue) (Fig. 2D). Subsequent Co-IP analyses demonstrated that both CBM1 and CBM2 independently bind Vago4 (Fig. 2E-F). *In vivo* treatment with exogenous recombinant CBM1 and CBM2 fragments significantly alleviated the inhibitory effect of Vago4 on EVC-induced STAT nuclear translocation compared to controls, as confirmed by immunofluorescence and Western blot analyses (Fig. 2G, g, H and h), indicating that these fragments are capable of competing with Domeless for binding to Vago4. These results confirm that Vago4 interacts with Domeless via the CBM1 and CBM2 domains.

**Figure 2.**
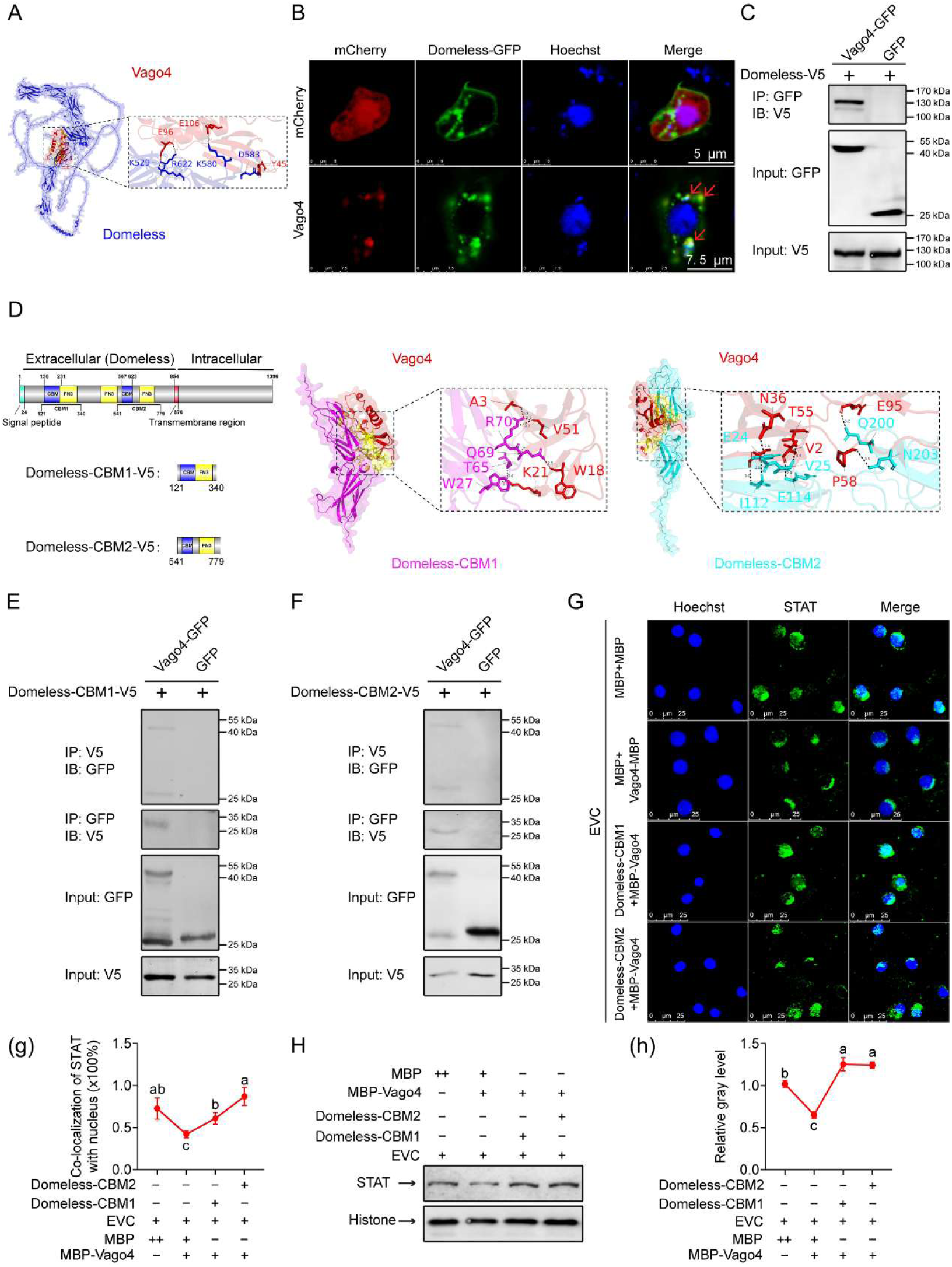
Interaction between Vago4 and Domeless. (A) Three-dimensional structures of the protein complexes formed by Vago4 with Domeless, predicted by AlphaFold3. (B) Immunofluorescence analysis showing the colocalization of Vago4 and Domeless in S2 cells. (C) Co-IP analysis confirming the interaction between Vago4 and Domeless. (D) Schematic representation of the domain structures of Domeless and its truncation mutants. Three-dimensional structures of the protein complexes formed by Vago4 with Domeless-CBM1 and Domeless-CBM2, respectively, predicted by AlphaFold3. (E–F) Co-IP analysis of the interactions between Vago4 and the CBM1 or CBM2 domains of Domeless. (G–H) Immunofluorescence and Western blot analyses of STAT nuclear translocation in hemocytes after *in vivo* co-injection of EVC with different recombinant protein combinations (MBP+MBP, Vago4-MBP+MBP, Vago4-MBP+Domeless-CBM1, or Vago4-MBP+Domeless-CBM2; 10 μg each) for 4 h. (g and h) Quantitative analyses of STAT–nucleus colocalization (panel G) and nuclear translocation proportion (panel H) using WCIF ImageJ software. Data are shown as mean ± SD. Statistical differences were determined by one-way ANOVA (g and h). Values with different letters (g and h) indicate significant differences. (*P* < 0.05).

### 2.3. Vago4 inhibits activation of JAK/STAT signaling by targeting Domeless

Immunofluorescence and Western blot analyses demonstrated that silencing of either *Vago4* or its upstream transcription factor *IRF* alone *in vivo* by RNAi strategy significantly promoted STAT nuclear translocation, with the effect being more pronounced following *Vago4* silencing (Fig. 3A, panels a, B, and b). Interestingly, the effect of simultaneously silencing *IRF* and *Vago4* was comparable to that of silencing *IRF* alone, but weaker than that of silencing Vago4 alone. This may be because *IRF* is also involved in other signaling pathways, and its silencing triggers certain feedback mechanisms that attenuate the effect of its downstream target gene *Vago4*, which requires further exploration. Previous studies have shown that STAT in shrimp inhibits the NF-κB family member Dorsal-mediated signaling pathway via a miRNA-1-mediated mechanism(Zuo *et al*., 2020). Based on this, we analyzed the expression of STAT and Dorsal downstream target genes in shrimp treated with recombinant Vago4 protein. The results showed that the expression of NF-κB downstream genes *ALF3*, *ALF4*, *PEN3*, and *LYZ2* was significantly upregulated upon Vago4 treatment, whereas the expression of the STAT downstream gene *ALF2* was markedly downregulated (Fig. 3C). Furthermore, the expression of the antimicrobial peptide gene *PEN4* was also significantly reduced upon Vago4 treatment (Fig. 3C). Using dual luciferase reporter gene assays, we further confirmed that the *PEN4* promoter is a downstream target of STAT (Fig. 3D). Expression of *STAT* or *Domeless* activated the *PEN4* promoter, and *Vago4* significantly attenuated the activating effect of Domeless. The *ie1* gene of WSSV can also be activated by STAT. Using the *ie1* promoter as a target, the dual-luciferase reporter gene assays further validated the effect of Vago4 on the Domeless receptor. *Vago4* expression significantly inhibited Domeless-mediated activation of the *ie1* promoter, and this inhibitory effect was weakened by mutation of the STAT-binding site within the *ie1* promoter (Fig. 3E).

**Figure 3.**
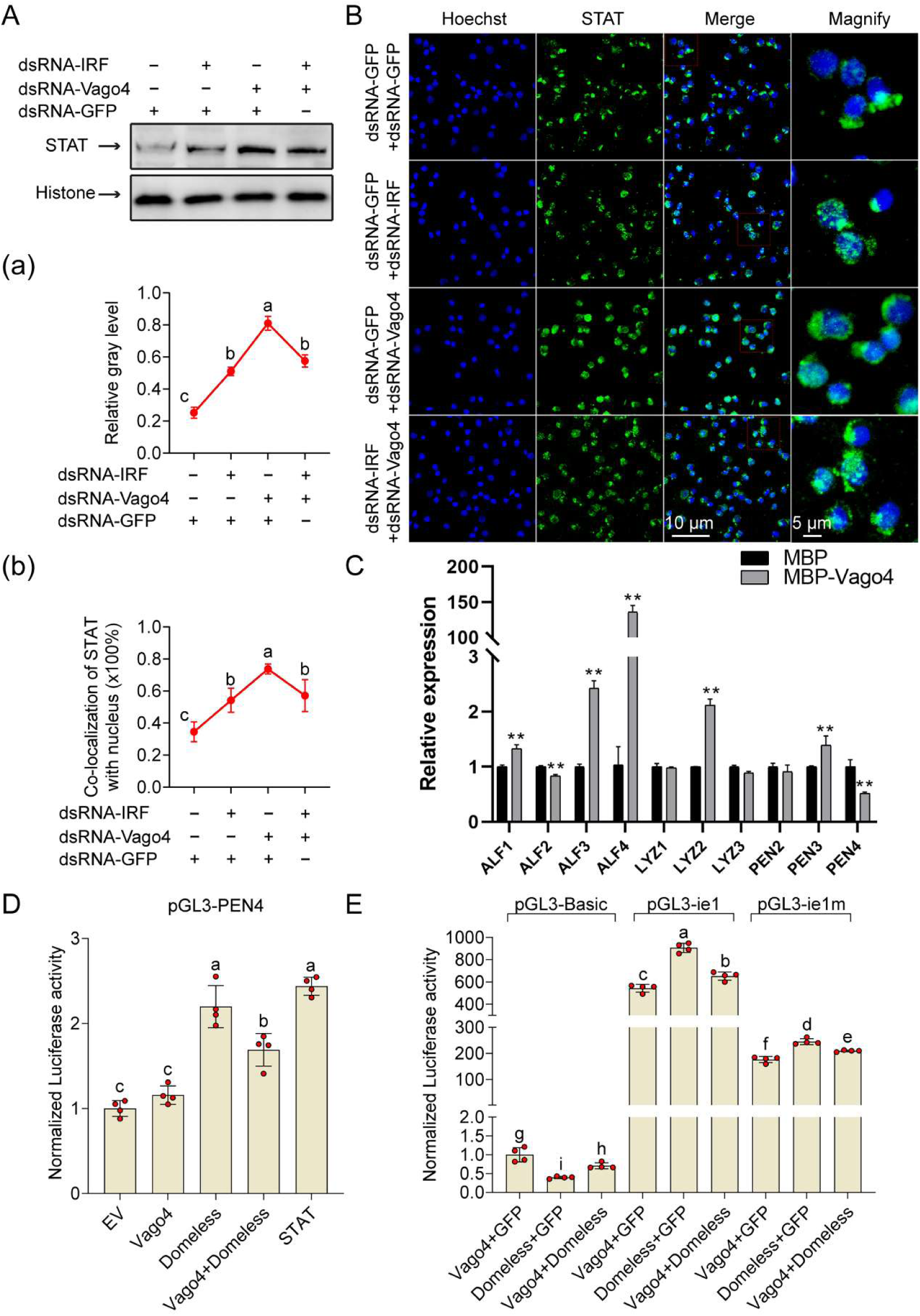
Vago4 suppresses Domeless-mediated activation of the JAK/STAT signaling pathway. (A) Western blot analysis of STAT nuclear translocation following *IRF* or *Vago4* silencing (48 h post 10 μg dsRNA/shrimp injection). (B) Immunofluorescence analysis of STAT nuclear localization after *IRF* or *Vago4* silencing. (C) qRT-PCR analysis of antimicrobial peptide gene expression following *in vivo* administration of recombinant Vago4 in shrimp. (D–E) Dual-luciferase reporter assays evaluating the effects of Domeless, Vago4, or STAT on the promoter activities of *PEN4*, *ie1*, and *ie1m* (STAT-binding site mutant). (a, b) Quantitative analysis of STAT–nucleus colocalization (panel A) and nuclear translocation proportion (panel B) using WCIF ImageJ software. The data are shown as the means ± SDs. Statistical differences were determined by one-way ANOVA (a-b, D and E) and Student *t*-test (C), **\*\*** *P*< 0.01. Values with different letters (a-b, D and E) indicate significant differences. (*P* < 0.05).

### 2.4. Vago4 induces ligand-dependent Domeless endocytosis and degradation

To investigate the mechanism of Vago4-mediated Domeless inhibition, we first examined its effect on Domeless protein abundance. Treatment with recombinant Vago4 markedly reduced Domeless protein levels in S2 cells, with maximal reduction observed 1 h after treatment (Fig. 4A). Increasing concentrations of recombinant Vago4 further decreased Domeless protein levels, indicating a dose-dependent effect (Fig. 4B). This effect was further confirmed *in vivo*, as the protein level of Domeless in shrimp hemocytes was significantly reduced following injection of recombinant Vago4 protein (Fig. 4C). Immunofluorescence analysis further showed that Vago4 treatment promoted redistribution of Domeless from the plasma membrane to the cytoplasm in hemocytes, indicating its internalization (Fig. 4D). To determine the mechanism underlying Domeless degradation, S2 cells were treated with the proteasome inhibitor MG132 and the lysosome inhibitor NH₄Cl. The results showed that both MG132 and NH4Cl blocked the Vago4-induced reduction in Domeless protein levels (Fig. 4E), and the inhibitory effect was concentration-dependent (Fig. 4F-G), suggesting that Domeless undergoes both proteasomal and lysosomal degradation upon internalization. Further analyses revealed that Vago4 promoted the polyubiquitination (Ub-HA) of Domeless (Fig. 4H). Analysis using single-lysine ubiquitin mutants further demonstrated that Vago4 specifically induced K27-linked ubiquitination of Domeless (Fig. 4I). In addition, exogenous Vago4 was internalized into lysosomes after uptake by S2 cells (Fig. 4J), while Vago4 treatment simultaneously promoted the trafficking of Domeless to lysosomes (Fig. 4K, k), suggesting that Vago4 and Domeless undergo coordinated trafficking to lysosomes following receptor internalization. To further validate these findings *in vivo*, shrimp were co-injected with recombinant Vago4 protein and pharmacological inhibitors. Western blot analysis showed that the Vago4-induced reduction in Domeless protein levels was partially reversed by either the lysosomal inhibitor chloroquine (CQ) or the clathrin-mediated endocytosis inhibitor chlorpromazine (CPZ) (Fig. 4L, l). Consistently, immunofluorescence analysis demonstrated that Vago4-induced redistribution of Domeless from the plasma membrane to the cytoplasm was markedly attenuated by CQ or CPZ treatment (Fig. 4N, Fig S3), indicating that both clathrin-mediated endocytosis and lysosomal trafficking are required for Vago4-induced Domeless internalization. Similarly, treatment with the proteasome inhibitor MG132 significantly restored Domeless protein levels (Fig. 4M, m), and immunofluorescence analysis further showed that MG132 partially reversed the Vago4-induced redistribution of Domeless from the plasma membrane to the cytoplasm in hemocytes (Fig. 4O, Fig S3), supporting a role for proteasomal degradation in Vago4-mediated Domeless downregulation following receptor internalization. Furthermore, STAT nuclear translocation was assessed *in vivo* to determine whether receptor endocytosis and degradation are required for Vago4-mediated inhibition of JAK/STAT signaling. Recombinant Vago4 markedly suppressed STAT nuclear translocation in hemocytes (Fig. 4P, p1, Fig S4). This inhibitory effect was partially reversed by CQ or CPZ (Fig. 4P, p2, Fig S4), indicating that receptor internalization is required for Vago4 function. Likewise, treatment with MG132 significantly restored STAT nuclear translocation (Fig. 4Q, q, Fig S4), demonstrating that proteasomal degradation of Domeless contributes to Vago4-mediated suppression of JAK/STAT signaling. Collectively, these findings demonstrate that Vago4 induces ligand-dependent Domeless endocytosis followed by coordinated lysosomal and K27-linked ubiquitination-dependent proteasomal degradation, thereby establishing receptor fate control as the mechanism underlying Vago4-mediated attenuation of JAK/STAT signaling.

**Figure 4.**
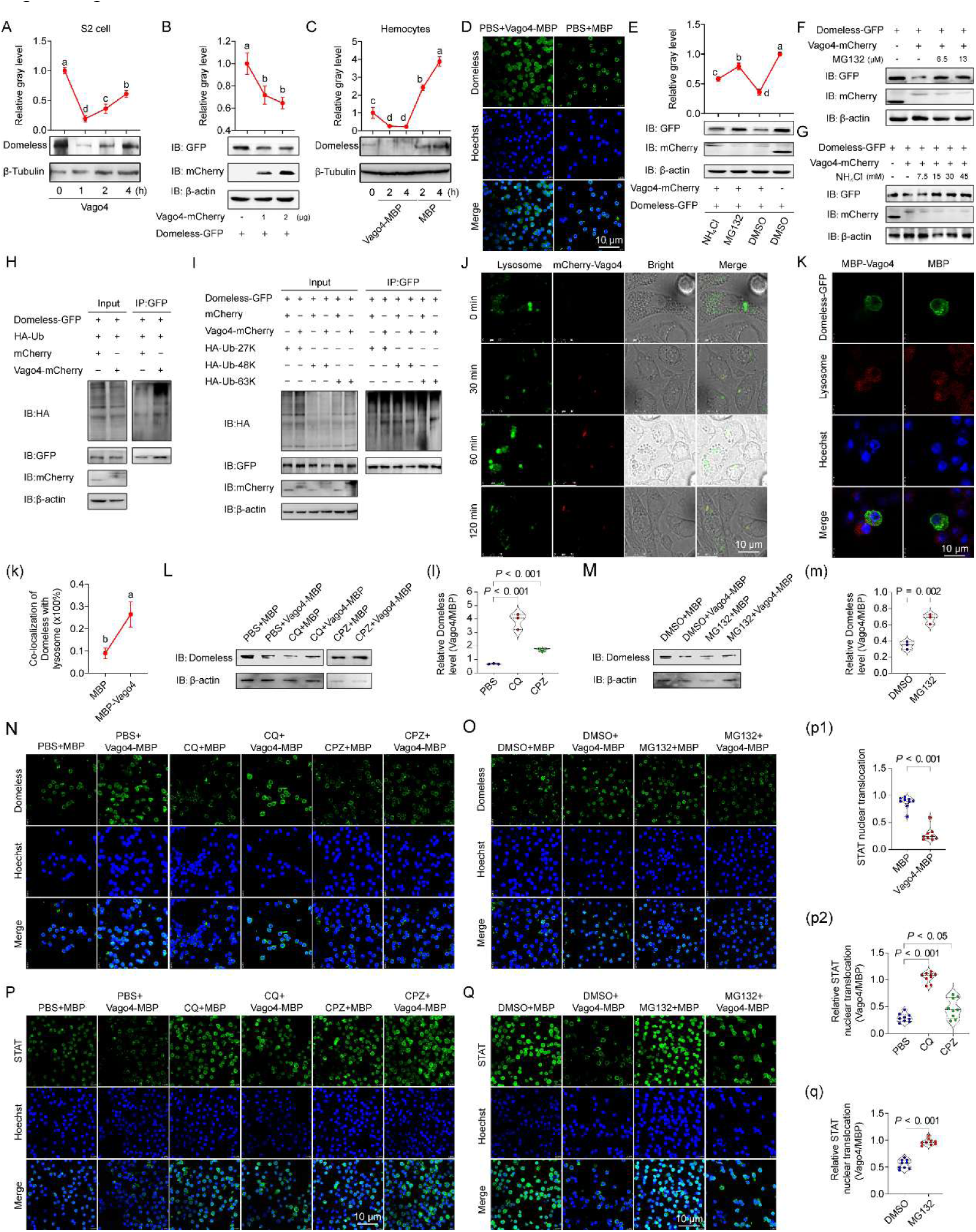
Vago4 induces ligand-dependent Domeless endocytosis and degradation. (A) Western blot analysis of Domeless protein levels following treatment with recombinant Vago4 (10 μg/ml, 1, 2 or 4 h) in S2 cells. Domeless protein levels were quantified by densitometric analysis using ImageJ. (B) Western blot analysis of Domeless protein levels in S2 cells after exposure to increasing concentrations of Vago4 protein (1 or 2 μg per well of a six-well plate). Domeless protein levels were quantified by densitometric analysis using ImageJ. (C-D) Western blot and immunofluorescence analyses of Domeless protein levels in shrimp following injection of recombinant Vago4 (10 μg). Western blot analysis was performed at 2 and 4 h post-injection, and immunofluorescence analysis was performed at 4 h post-injection. Domeless protein levels were quantified by densitometric analysis using ImageJ. (E) Western blot analysis of Domeless protein levels in S2 cells treated with Vago4 in the presence of the proteasome inhibitor MG132 (6.5 μM, 4 h) or the lysosomal inhibitor NH₄Cl (7.5 mM, 4 h). Domeless protein levels were quantified by densitometric analysis using ImageJ. (F-G) Western blot analysis of Domeless protein levels following 4 h treatment with graded concentrations of MG132 (6.5 or 13 μM) or NH₄Cl (7.5 to 45 μM) in the presence of Vago4. (H) Western blot analysis of Vago4-induced changes in Domeless ubiquitination levels (Ub). (I) Western blot analysis of Domeless ubiquitination in the presence of Vago4 and ubiquitin mutants (Ub-K27, Ub-K48, and Ub-K63). (J) Exogenous mCherry-tagged Vago (red, expressed and purified from S2 cells) was localized to lysosomes (stained with Lyso-Tracker Green) in S2 cells within 1–120 min of treatment (10 μg/mL). (K) Recombinant Vago4 protein treatment (10 μg/mL, 2 h) induced colocalization of GFP-tagged Domeless with lysosomes (stained with Lyso-Tracker Red) in S2 cells. Statistical analysis of the colocalization ratio was performed using ImageJ. (L, N) Western blot (L) and immunofluorescence (N) analysis of Domeless protein levels in shrimp hemocytes 4 h after co-injection of recombinant Vago4 protein (10 μg/shrimp) with either chloroquine (CQ, 100 μg/shrimp), chlorpromazine (CPZ, 50 μg/shrimp) or PBS. Domeless protein levels were quantified by densitometric analysis using ImageJ (l). (M, O) Western blot (M) and immunofluorescence (O) analysis of Domeless protein levels in shrimp hemocytes 4 h after co-injection of recombinant Vago4 protein (10 μg/shrimp) with either MG132 (20 μg/shrimp), or DMSO. Domeless protein levels were quantified by densitometric analysis using ImageJ (m). (P, p) Immunofluorescence analysis of STAT nuclear translocation in hemocytes 4 h after co-injection of recombinant Vago4 protein (10 μg) with either chloroquine (CQ, 100 μg), chlorpromazine (CPZ, 50 μg) or PBS. STAT nuclear translocation was quantified by ImageJ. Data are presented STAT nuclear translocation in Vago4-treated samples compared with the corresponding MBP control (p). Data are presented as the fold change of Vago4-treated samples relative to the corresponding MBP control (PBS+Vago4-MBP/PBS-MBP, CPZ+Vago4-MBP/CPZ-MBP, or CQ+Vago4-MBP/CQ-MBP) (p1-2). (Q, q) Immunofluorescence analysis of STAT nuclear translocation in hemocytes 4 h after co-injection of recombinant Vago4 protein (10 μg) with MG132 (20 μg) or DMSO control. STAT nuclear translocation was quantified by ImageJ. Data are presented as the fold change of Vago4-treated samples relative to the corresponding MBP control (DMSO+Vago4-MBP/DMSO-MBP or MG132+Vago4-MBP/MG132-MBP) (q). Data are shown as the mean ± SD. Statistical differences were determined by one-way ANOVA (A-C, E, k, l, m, p and q). Values with different letters (A-C, E and k) indicate significant differences. (*P* < 0.05).

### 2.5. The role of Vago4–JAK/STAT signaling in acute V. parahaemolyticus infection

Previous studies have demonstrated that acute *V. parahaemolyticus* infection induces excessive activation of the JAK/STAT pathway in shrimp, resulting in characteristic inflammatory lesions, including opaque white regions at the intramuscular injection site, widened intermuscular spaces, and hemocyte infiltration(Zuo *et al*., 2025). This response is accompanied by increased reactive oxygen species (ROS) levels in hemocytes and impaired phagocytic activity(Zuo *et al*., 2025). To investigate the role of Vago4 during this process, shrimp were subjected to *in vivo* silencing of *Vago4* (Fig. 5A), followed by acute infection with 1.5 × 10⁶ CFU of *V. parahaemolyticus*. *Vago4* silencing significantly decreased shrimp survival rates (Fig. 5B). Consistent with this, injection or feeding with recombinant Vago4 protein significantly improved post-infection survival (Fig. 5C-D). These suggest that Vago4 plays a protective role in host defense during acute infection.

**Figure 5.**
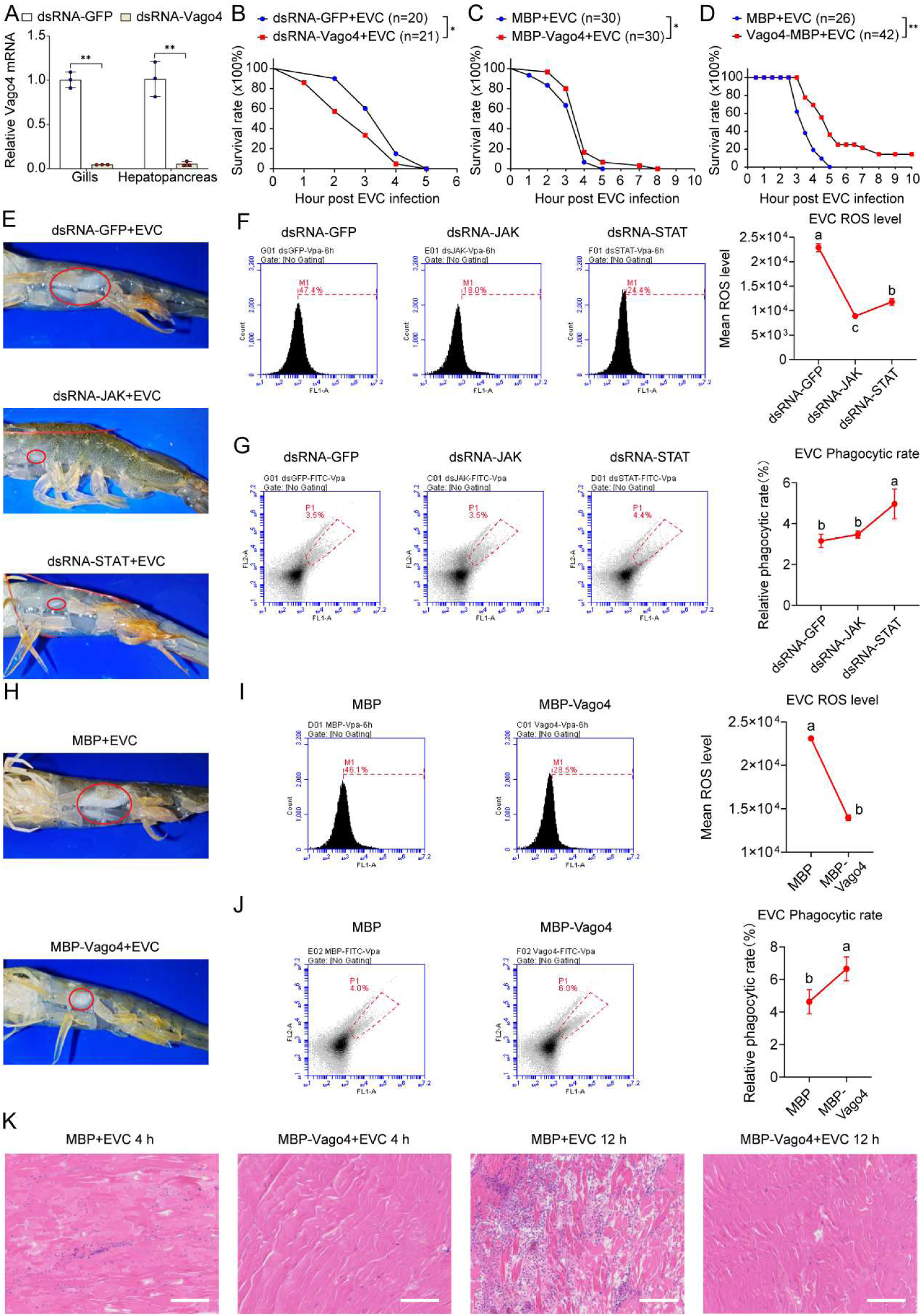
The role of the Vago4–JAK/STAT pathway in acute *V. parahaemolyticus* infection. (A) qRT-PCR analysis of *Vago4* RNAi knockdown efficiency (10 μg dsRNA/shrimp, 48 h) in gills and hepatopancreas. (B) Survival analysis of shrimp treated with dsRNA (10 μg, 48 h) following infection with *V. parahaemolyticus* (1.5 × 10⁶ CFU). * *P* < 0.05 by log-rank (Mantel-Cox) test. (C-D) Survival analysis of shrimp injected (C) or fed (D, 0.125 mg per shrimp per day, for seven days) recombinant Vago4 protein following infection with *V. parahaemolyticus* (1.5 × 10⁶ CFU). * *P* < 0.05 by log-rank (Mantel-Cox) test. (E-J) The pathological appearance (E and H), hemocyte ROS level (F and I) and phagocytic activity (G and J) following knockdown of *JAK*, *STAT* (10 μg dsRNA/shrimp, 48 h), or Vago4 proteins (10 μg/shrimp, 4 h) treatment and subsequent infection with a high dose of *V. parahaemolyticus* (1.5 × 10⁶ CFU). Opaque whitish lesions in muscle tissue are indicated by red circles. Hemocytes were collected from nine shrimp per experimental group. (K) The hematoxylin–eosin (H&E) staining of shrimp muscle tissue following Vago4 protein treatment and subsequent infection with a high dose of *V. parahaemolyticus* (1.5 × 10⁶ CFU). Data are shown as means ± SD. Statistical differences were determined by one-way ANOVA (F, G, I and J). Values with different letters (F, G, I and J) indicate significant differences. (*P* < 0.05).

To further assess whether Vago4 function depends on the JAK/STAT pathway, key downstream components, JAK and STAT, were individually silenced. Compared with controls, silencing of either *JAK* or *STAT* in acute infected shrimp reduced the formation of opaque white lesions at the intramuscular injection site (Fig. 5E) and decreased reactive oxygen species (ROS) levels in hemocytes (Fig. 5F). Similarly, recombinant Vago4 treatment alleviated this inflammatory-like phenotype (Fig. 5H) and reduced ROS levels in hemocytes (Fig. 5I). Previous studies have shown that silencing of *JAK* and *STAT* under acute infection conditions enhances the phagocytic activity of hemocytes(Zuo *et al*., 2025). This observation was further confirmed in the present study (Fig. 5G). Likewise, treatment with recombinant Vago4 protein also significantly increased hemocyte phagocytic capacity (Fig. 5J), providing additional evidence that Vago4 suppresses JAK–STAT pathway activation during acute infection. Histological analysis further revealed that recombinant Vago4 attenuated myofiber gap widening and hemocyte infiltration at both 6 h and 12 h post-infection (Fig. 5K). Collectively, these results demonstrate that Vago4 exerts protection against acute *V. parahaemolyticus* infection by inhibiting the JAK/STAT pathway. This mechanism may be associated with reduced oxidative stress and enhanced phagocytosis in hemocytes, thereby mitigating inflammatory tissue damage in shrimp.

## 3. Discussion

It is now well established that both invertebrates and vertebrates possess cytokine-mediated immune signaling pathways linking IRF to JAK/STAT signaling(Hou Y *et al*., 2025; Li *et al*., 2015; Paradkar *et al*., 2012; Stark and Darnell, 2012). In arthropods, the Vago-mediated signaling axis is regarded as a key immune regulatory module functionally analogous to the vertebrate IFN system and has attracted increasing attention in recent years(Hou Y *et al*., 2025; Li *et al*., 2015; Paradkar *et al*., 2012). Although Vago shares no sequence homology with mammalian IFNs, the antiviral immunity it mediates exhibits IFN-like functional features(Gao *et al*., 2021; Li *et al*., 2015; Paradkar *et al*., 2012). On one hand, Vago is induced by the viral dsRNA sensor Dicer-2, whose recognition pattern shows partial conservation with mammalian RIG-I-like receptors. On the other hand, Vago in mosquitoes and shrimp elicits antiviral responses through activation of the JAK/STAT pathway, mirroring the functional strategy of vertebrate IFNs(Deddouche *et al*., 2008; Gao *et al*., 2021; Li *et al*., 2015; Paradkar *et al*., 2012). Therefore, these observations support the view that the Vago signaling axis constitutes an invertebrate immune regulatory module functionally corresponding to the IFN system. It is worth noting that, although invertebrate Domeless, as a conserved homolog of Class I cytokine receptors, is known to mediate JAK/STAT signaling, there is currently a lack of evidence regarding whether Vago acts as a functional ligand to directly regulate Domeless(Sun *et al*., 2017). Here, we identify Vago4 as a bona fide ligand of Domeless and demonstrate that receptor recognition is mediated through the extracellular CBM1 and CBM2 domains of Domeless. These findings further refine the IRF–Vago–Domeless–JAK/STAT IFN-like signaling axis and strengthen its functional analogy to the vertebrate IFN system.

Excessive activation of immune signaling pathways can convert protective immune responses into pathogenic processes, leading to immunopathological damage and even host death; thus, restricting excessive signal amplification and maintaining immune homeostasis are essential for effective host defense(Medzhitov R *et al*., 2012; Zuo *et al*., 2025). Endocytosis and degradation of cell surface receptors represent key negative regulatory mechanisms that limit signal intensity and duration(Bhattacharya *et al*., 2010; Marijanovic *et al*., 2006). Multiple immune receptors, including TLR4, EGFR, and TCR, undergo ligand-induced endocytosis coupled with coordinated proteasomal and lysosomal degradation, thereby reducing receptor abundance and limiting signal transduction(Balagopalan *et al*., 2009; Husebye *et al*., 2006; Levkowitz *et al*., 1999). Under immune hyperactivation, cytokine-driven receptor downregulation represents a critical negative-feedback mechanism that limits sustained signal amplification(Bhattacharya *et al*., 2010; Marijanovic *et al*., 2006). In the IFN system, for example, IFNα induces phosphorylation, ubiquitination, and degradation of IFNAR1, thereby limiting the sustained activation of the JAK/STAT pathway(Bhattacharya *et al*., 2010). Similar receptor downregulation mechanisms have also been described in inflammatory signaling pathways such as TNFR(Wajant and Scheurich, 2011). Although these mechanisms have been extensively elucidated in mammals, whether invertebrate cytokines similarly suppress immune overactivation by regulating receptor fate remains largely unknown. As the only type I cytokine receptor identified in invertebrates, the endocytic degradation of Domeless was initially observed in *Drosophila*, but the underlying mechanism remained unclear(Hou *et al*., 2015; Vidal *et al*., 2010). Here, we demonstrate that Vago4 initiates Domeless endocytosis and receptor downregulation, thereby restraining excessive JAK/STAT activation during acute infection. Pharmacological inhibition of clathrin-mediated endocytosis or receptor degradation largely abolished the inhibitory effect of Vago4 on STAT activation, demonstrating that receptor fate control is essential for Vago4-mediated immune suppression. Functionally, this resembles the regulatory pattern observed in vertebrates, where cytokines such as IFN-γ not only initiate anti-infective responses but also contribute to limiting excessive inflammation(Blouin *et al*., 2016; Hu and Ivashkiv, 2009; Ivashkiv, 2018). These findings uncover a ligand-induced receptor downregulation mechanism that maintains immune homeostasis in invertebrates and further highlight the dual roles of Vago4 in both immune activation and immune suppression.

Ubiquitination-mediated protein degradation plays a pivotal role in the dynamic regulation of cell surface receptors(Guo T *et al*., 2019). Following ligand stimulation, receptors are typically modified by E3 ubiquitin ligases, marking them for endocytic trafficking and subsequent degradation(Haglund and Dikic, 2012; Husebye *et al*., 2006). Although the degradation of membrane receptors primarily relies on the lysosomal pathway, ubiquitination serves not only as a degradation signal but also as a crucial marker for endocytic sorting, determining the receptor’s transport pathway and ultimate fate(Haglund and Dikic, 2012). In the regulation of immune signaling, the ubiquitin-dependent degradation mechanism is crucial for limiting sustained signal activation and preventing immune overreactions(Haglund and Dikic, 2012; Husebye *et al*., 2006). Here, we identify K27-linked ubiquitination as a key molecular event coupling Domeless endocytosis to coordinated lysosomal and proteasomal degradation, thereby restraining excessive JAK/STAT activation. The specific E3 ubiquitin ligase responsible for Vago4-induced Domeless K27-linked ubiquitination and degradation requires further identification(Zhang W *et al*., 2023). This will help refine the mechanism underlying Vago4-mediated receptor downregulation and deepen our understanding on the fine-tuned regulatory network of JAK/STAT signaling.

Administration of recombinant Vago4 protein alleviated inflammatory symptoms and reduced mortality in shrimp acutely infected with *V. parahaemolyticus*. These findings suggest that Vago4-mediated receptor downregulation provides a potential strategy for precisely controlling excessive JAK/STAT activation. Therefore, developing targeted immunomodulatory strategies directed at the Vago4-Domeless axis, such as utilization of recombinant Vago4 proteins, functional peptides, or gene expression regulatory tools, may offer promising approaches to mitigate excessive immune responses, alleviate tissue damage during acute infection, and ultimately enhance shrimp survival and improve the stability of aquaculture production. Moreover, identifying or engineering key molecular regulators of Domeless endocytosis and degradation, including ubiquitin-associated enzymes and endocytic regulators, may facilitate the fine-tuning of JAK/STAT signal intensity and duration. Furthermore, elucidating whether Vago4 acts synergistically with other immune modulators to maintain immune homeostasis could provide a theoretical basis for developing multi-target intervention strategies. Beyond shrimp aquaculture, these mechanistic insights may extend to other invertebrates and aquatic host–pathogen systems, offering a broader framework for understanding conserved principles for maintaining immune homeostasis. Although this study establishes a critical role for Vago4 in restraining excessive JAK/STAT activation, several important questions remain unresolved. For example, whether Vago4-mediated receptor degradation influences crosstalk with other signaling pathways, and how Vago4 is dynamically regulated across different stages of infection, warrant further investigation. Addressing these questions will advance our understanding of how ligand-induced receptor fate control shapes immune homeostasis and cytokine signaling in invertebrates, while providing broader insights into the evolution and fine-tuning of JAK/STAT signaling.

In summary, this study identifies Vago4 as a bona fide ligand of the canonical JAK/STAT receptor Domeless and demonstrates that Vago4 induces Domeless endocytosis, coupled with K27-linked ubiquitination-dependent lysosomal and proteasomal degradation, thereby restraining excessive JAK/STAT activation (Fig. 6). These findings provide direct molecular evidence for cytokine-mediated receptor downregulation in invertebrates, identify ligand-induced receptor fate control as a previously unrecognized negative-feedback mechanism for maintaining immune homeostasis, and provide new insights into the evolution of interferon-like cytokine signaling. Together, our findings extend the functional paradigm of interferon-like cytokines from signal initiation to dynamic receptor fate control, revealing a conserved strategy for maintaining immune homeostasis in invertebrates.

**Figure 6.**
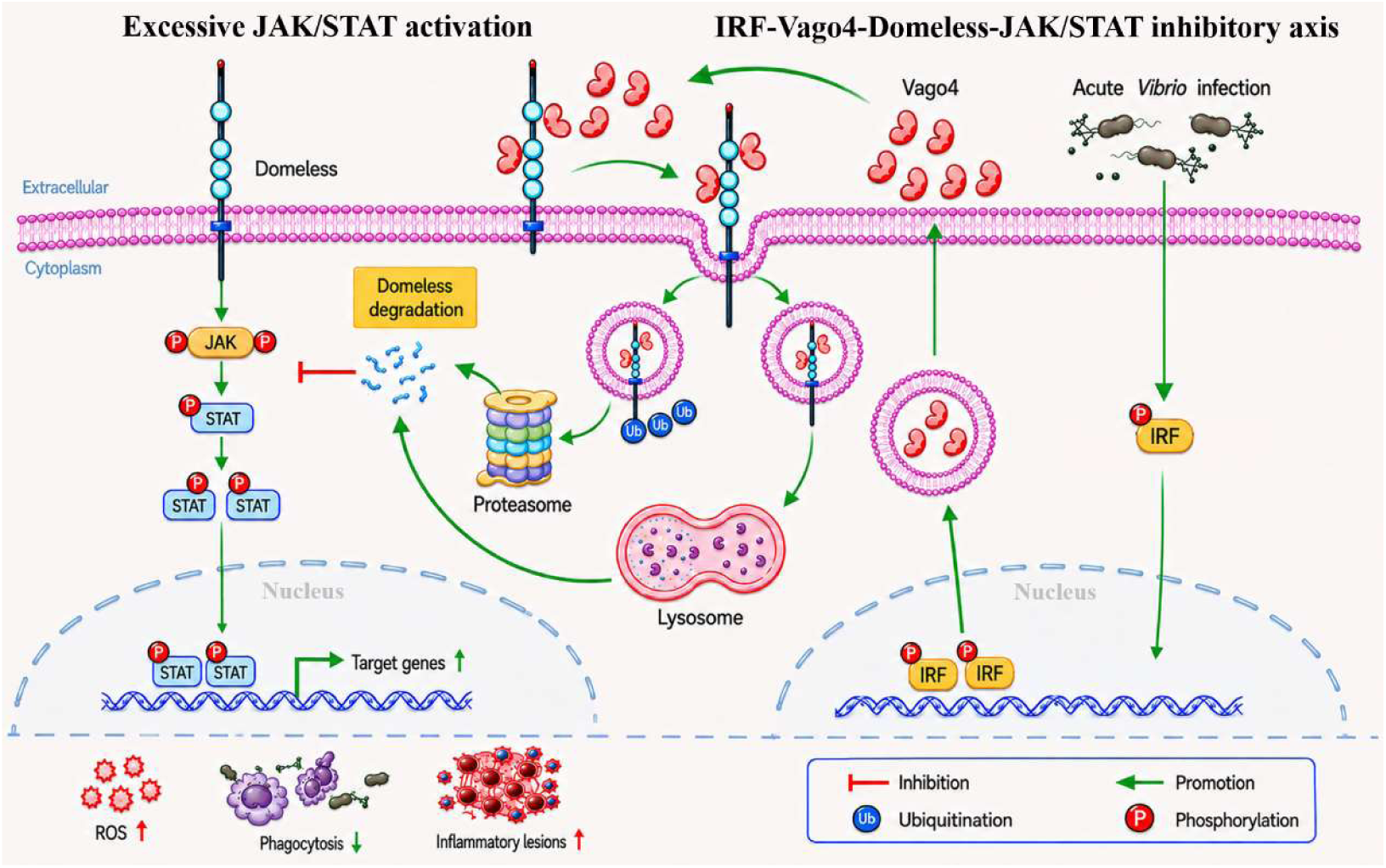
Proposed model of Vago4-mediated negative regulation of Domeless–JAK/STAT signaling during acute *Vibrio* infection. Acute *Vibrio* infection activates JAK/STAT signaling through Domeless, resulting in STAT phosphorylation and transcriptional activation of target genes. Excessive pathway activation is associated with increased ROS production, impaired phagocytosis and inflammatory tissue damage. Infection-induced IRF activation promotes Vago4 transcription, and secreted Vago4 binds Domeless and triggers receptor endocytosis, followed by lysosomal trafficking and ubiquitination-dependent proteasomal degradation. Domeless downregulation consequently attenuates JAK/STAT signaling and restrains excessive immune activation, thereby reducing ROS production and inflammatory damage while enhancing phagocytic activity and host survival. Green arrows indicate promotion, red T-bars indicate inhibition, red circles labeled “P” indicate phosphorylation, and blue circles labeled “Ub” indicate ubiquitination.

## 4. Materials and Methods

### Animal and pathogen

Healthy *P. vannamei* (approximately 10 g) were obtained from Foshan, China, and acclimated for at least one week in a recirculating aquaculture system containing aerated water with a salinity of 5 ppt at 27 °C. The system is equipped with an ultraviolet sterilization filter for continuous removal of pathogenic microorganisms. Shrimp were fed a commercial artificial diet daily at 3% body weight, divided into two feeding periods. The *V. parahaemolyticus* stored in our lab(Xiao B *et al*., 2023) was cultured in LB and quantified using the plate counting method.

### RNAi and qRT-PCR

RNA interference (RNAi) was employed to suppress expression of the *IRF*, *Vago4*, *JAK*, and STAT genes. dsRNAs were synthesized using the T7 RiboMAX™ Express RNAi System (Promega, USA). Briefly, gene-specific templates were amplified using primers containing the T7 polymerase promoter sequence (Table S1). Complementary single-stranded RNAs were synthesized separately and subsequently annealed to generate dsRNA. dsRNA targeting *GFP* gene was used as a control. For gene silencing, 10 μg of dsRNA in 50 μL PBS was injected into the second abdominal segment of shrimp. At 48 h post-injection, the mRNA levels of *IRF*, *Vago4*, *JAK*, and *STAT* in the gill, and hepatopancreas tissues were quantified by qRT-PCR to evaluate RNAi efficiency and gene expression.

For *in vivo* RNAi, shrimp were divided into two groups (n = 20–21) and injected with Vago4-specific dsRNA or control GFP dsRNA, as described above. After 48 h, shrimp were challenged with 1.5 × 10^6^ CFU of *V. parahaemolyticus*, and mortality was monitored.

For acute infection following JAK/STAT pathway downregulation, shrimp were divided into three groups (n = 30) and injected with dsRNA targeting *JAK*, *STAT*, or control *GFP*, respectively, as described above. After 48 h, shrimp were challenged with 1.5 × 10^6^ CFU of *V. parahaemolyticus*. At 4 hpi, hemocytes were collected from nine shrimp per group, and hemocytes were collected for flow cytometry analysis.

Total RNA was extracted and reverse-transcribed into cDNA using the PrimeScript RT Kit with gDNA Eraser (Takara, Japan). Each qRT-PCR reaction was performed in a final volume of 10 μL containing 1 μL cDNA, 5 μL 2× SYBR Premix Ex Taq II (Takara, Japan), and 500 nM of each gene-specific primer (Table S1) using a LightCycler 480 System (Roche, Switzerland) under the following conditions: initial denaturation at 95°C for 1 min, followed by 40 cycles of denaturation at 95°C for 10 s and annealing/extension at 60°C for 30 s. Relative gene expression levels were calculated using the 2^−ΔΔCt^ method, with elongation factor 1α (*EF-1α*; GenBank accession no. GU136229) used as the internal reference gene. All experiments were independently repeated using three different batches of shrimp, and representative results are presented.

### Recombinant expression and purification

The *Vago4* gene lacking the signal peptide-coding sequence was cloned into the pMAL-c2X vector. The recombinant plasmid pMAL-c2X-*Vago4* and the empty pMAL-c2X vector were individually transformed into *E. coli* Rosetta (DE3) cells for MBP-tagged Vago4 and MBP expression. Recombinant proteins were purified using amylose resin (New England Biolabs, USA) according to the manufacturer’s instructions. Gene fragments encoding the extracellular CBM1 and CBM2 domains of Domeless were cloned into the pET-32a (+) vector (Merck, Germany). The recombinant plasmids, together with the empty pET-32a (+) vector, were separately transformed into *E. coli* BL21(DE3) competent cells (Invitrogen, USA). Following IPTG induction, recombinant proteins were purified using Ni-NTA agarose resin (Qiagen, Germany) according to the manufacturer’s instructions. The mCherry-tagged Vago4 protein was purified from the supernatant of S2 cell cultures as previously described(He J *et al*., 2023).

### Immune challenge

In the immune challenge experiments were independently repeated three times, and the collected samples were subsequently subjected to qRT-PCR, Western blot, flow cytometry, and immunofluorescence analyses. For immune challenge, shrimp were randomly divided into four groups (n = 30) and intramuscularly injected with 50 μL PBS (as control), or 50 μL PBS containing 5 × 10^5^ CFU (NVC), or 1.5 × 10^6^ CFU (EVC) of *V. parahaemolyticus*, or 5 μg poly(I:C). At each time points post injection, hemocytes were collected from nine shrimp per group, and hemocyte smears were prepared for immunofluorescence analysis.

### Protein treatment experiments

For *in vitro* experiments, *Drosophila* S2 cells ectopically expressing shrimp STAT were incubated with recombinant Vago4-MBP or MBP control protein at final concentrations of 1, 5, or 10 μg/mL together with poly(I:C) (0.5 μg/mL) for 4 h. Cells were then harvested for Western blot and immunofluorescence analyses. For *in vivo* experiments, shrimp were randomly assigned to six groups (n = 30) and individually injected with *V. parahaemolyticus* (1.5 × 10⁶ CFU in 50 μL PBS) plus one of the following supplements: MBP (10 μg), Vago4-MBP (10 μg), MBP + MBP (10 μg each), Vago4-MBP + MBP (10 μg each), Vago4-MBP + Domeless-CBM1 (10 μg each), or Vago4-MBP + Domeless-CBM2 (10 μg each). After 4 h, hemocytes were collected from nine shrimp per group for Western blot, immunofluorescence, and flow cytometry analyses. The muscle tissues at the injection site of *V. parahaemolyticus* from Vago4-MBP/MBP-treated shrimp were collected, fixed, paraffin-embedded, sectioned, and subjected to HE staining histopathological analysis. In parallel experiments, the mortality of *V. parahaemolyticus*-infected shrimp was recorded.

### Co-immunoprecipitation (Co-IP), Western blot and nuclear-cytoplasmic separation

S2 cells were co-transfected with the indicated plasmids (pAc5.1 (invitrogen) expression vectors encoding *P. vannamei* Vago4-GFP, GFP, Domeless-CBM1-V5 and Domeless-CBM2-V5) and harvested at 48 h post-transfection. Cells were lysed for 30 min at 4 °C using lysis buffer (#P0013, Beyotime, China) supplemented with a protease inhibitor cocktail (#P8340, Thermo Fisher Scientific, USA), followed by centrifugation at 13,000 × g for 5 min. The resulting supernatants were collected for Co-IP using magnetic beads conjugated with the indicated antibodies according to the manufacturer’s instructions. For Co-IP assays, anti-GFP antibody (#3057, Dia-An Biotech), anti-V5 antibody (#AB3792, Merck Millipore), or anti-HA antibody (3063R, Dia-An Biotech) was used. Proteins obtained from Co-IP or total cellular proteins prepared were treated with SDS loading buffer and separated on 10% SDS-PAGE and transferred onto PVDF membranes (Millipore, USA). Membranes were blocked 5% non-fat milk and incubated with the indicated primary antibodies. The following primary antibodies were used for Western blot: anti-GFP antibody (#3057, Dia-An Biotech), anti-V5 antibody (#AB3792, Merck Millipore), anti-phospho-STAT antibody (customized from GL Biochem, Shanghai, China), anti-shrimp STAT antibody (customized from GL Biochem, Shanghai, China), anti-actin clone C4 Abs (#MAB1501, Merck Millipore), and anti-Histone H3 antibody (#4499S, CST). After three washes, membranes were incubated with HRP-conjugated goat anti-rabbit IgG (#w4011, Promega) or goat anti-mouse IgG (#w4021, Promega) for 1 h at room temperature. Protein bands were visualized using a chemiluminescence imaging system (#5200, Tanon, China), and band intensities were quantified using ImageJ software. For STAT activation assays, nuclear and cytoplasmic fractions were separated using a nuclear and cytoplasmic extraction kit (NE-PER™, #78833, Thermo Fisher Scientific) according to the manufacturer’s instructions, followed by Western blot analysis of each fraction.

### Ubiquitination Assays

The GFP-tagged Domeless plasmid, wild-type ubiquitin (Ub-HA), ubiquitin mutants (K27-HA, K48-HA, or K63-HA), and empty vector (EV, for mCherry expression) were co-transfected into *Drosophila* S2 cells together with the Vago4-mCherry expression vector. At 36 h post-transfection, cells were treated with either the proteasome inhibitor MG132 (13 μM; #S2619, Selleck, USA) or an equivalent volume of DMSO for 4 h. Cell lysates were subsequently subjected to immunoprecipitation using anti-GFP magnetic beads, followed by Western blot analysis with the indicated antibodies.

### Flow cytometry

Flow cytometry analysis was performed to evaluate intracellular reactive oxygen species (ROS) production and phagocytic capacity in hemocytes derived from shrimp subjected to STAT or JAK knockdown. For each treatment group, hemocytes were obtained from nine individuals, and the collected hemocytes were rinsed three times with 2× Leibovitz’s L-15 medium (Gibco, USA) before further processing.

For ROS measurement, hemocytes were incubated with the fluorescent probe 2′,7′-dichlorodihydrofluorescein diacetate (DCFH-DA) at a working concentration of 10 μmol/L for 20 min at 37 °C under dark conditions. Following incubation, the cells were washed thoroughly, and intracellular fluorescence intensity was quantified using a C6 flow cytometer (BD Biosciences, USA).

For phagocytosis assay, FITC-labeled *V. parahaemolyticus* were prepared as previously described(Zuo *et al*., 2025). Hemocytes were stained with DiI membrane dye (Beyotime, China) and subsequently co-incubated with FITC-labeled *V. parahaemolyticus* at a hemocyte-to-bacterium ratio of 1:100 for 1 h at 28 °C. After washing three times with L-15 medium, cells were subjected to flow cytometry to analyze fluorescence signals of DiI (Ex/Em = 549/565 nm) and FITC (Ex/Em = 488/519 nm). The phagocytosis-positive gates were defined by analyzing DiI-labeled hemocytes incubated with unlabelled bacteria and unstained hemocytes incubated with FITC-labeled bacteria. For each sample, 100,000 events were analyzed.

### Immunofluorescence

S2 cells were cultured on silane-treated coverslips placed in 24-well culture plates and transfected with plasmids expressing the indicated fluorescently tagged proteins. At 36 h post-transfection, cells were fixed with 4% paraformaldehyde and counterstained with Hoechst 33258. Fluorescence signals were subsequently visualized using a Leica TCS SP8 confocal laser-scanning microscope (Germany).

Hemocyte samples were spread onto silane-coated glass slides and fixed in 4% paraformaldehyde for 10 min, followed by membrane permeabilization with 1% Triton X-100 for 20 min. The slides were then incubated with rabbit polyclonal antibodies against shrimp STAT protein (GL Biochem, China), followed by Alexa Fluor 488-conjugated goat anti-rabbit IgG (Abcam, USA). Nuclear staining was subsequently performed using Hoechst 33342 (Invitrogen, USA). Fluorescence signals were subsequently visualized using a Leica TCS SP8 confocal laser-scanning microscope (Germany).

For inhibitor treatment, shrimp were co-injected with recombinant Vago4 protein (10 μg) and either PBS (control), the clathrin-mediated endocytosis inhibitor chlorpromazine (CPZ, 50 μg; #A506232, Sangon, China), the lysosomal inhibitor chloroquine (CQ, 100 μg; #HY-17589A, MCE, USA), or the proteasome inhibitor MG132 (20 μg; #S2619, Selleck, USA). Hemocytes were collected 4 h after injection for immunofluorescence analysis.

For lysosomal localization assays, *Drosophila* S2 cells were incubated with recombinant mCherry-tagged Vago4 protein (10 μg/mL), which was expressed and purified from S2 cells, for the indicated times (0, 30, 60, or 120 min). Cells were subsequently stained with LysoTracker Green (#L7526, Thermo Fisher Scientific) according to the manufacturer’s instructions, fixed with 4% paraformaldehyde for 15 min at room temperature, and imaged using a Leica TCS SP8 confocal microscope (Germany).

For analysis of Domeless trafficking, S2 cells expressing GFP-tagged Domeless were treated with recombinant Vago4 protein (10 μg/mL) for 4 h, followed by staining with LysoTracker Red (#L7528, Thermo Fisher Scientific) according to the manufacturer’s instructions. Cells were fixed with 4% paraformaldehyde for 15 min at room temperature and imaged using the same confocal microscope. Colocalization between Domeless and lysosomes was quantified using the JACoP plugin in ImageJ, and the colocalization ratio was calculated from at least three independent experiments.

### Luciferase Reporter Assay

To evaluate the regulatory effect of Vago4 on Domeless-mediated signaling, pAc5.1 (Invitrogen, USA) expression vectors encoding *P. vannamei* Vago4, Domeless, and STAT, and the pGL3-Basic reporter vectors (Promega, USA) containing the *PEN4*, *ie1*, or STAT-binding motif-mutated *ie1* (*ie1m*) promoters were used(Yan *et al*., 2015). S2 cells were seeded in 12-well culture plates and transfected with expression plasmids, including Vago4 alone, Domeless alone, co-combination of Vago4 and Domeless, or STAT alone (as positive control) together with corresponding firefly luciferase reporter plasmids and the pRL-TK Renilla luciferase vector (as internal control, Promega, USA). At 48 h post transfection, cell lysates were collected from six independent biological replicates per group, and luciferase activities were quantified using a Dual-Luciferase Reporter Assay System (Promega, USA). Relative promoter activity was calculated based on the ratio of firefly to Renilla luciferase activity.

## Statistical Analysis

Differences between two groups were evaluated using an unpaired two-sided Student’s *t*-test, and comparisons involving multiple groups were analyzed by one-way ANOVA followed by Dunnett’s post *hoc* test. Survival curves were assessed using Kaplan–Meier survival analysis in conjunction with the log-rank χ2 test. Statistical significance was defined as *P* < 0.05.

## 5. Author Contributions

**Conceptualization**: Xiaopeng Xu, Jianguo He.

**Formal analysis**: Linwei Yang.

**Funding acquisition**: Linwei Yang, Hongliang Zuo, Jianguo He, Xiaopeng Xu.

**Investigation**: Linwei Yang, Jianhui He.

**Methodology**: Xiaopeng Xu.

**Data curation**: Linwei Yang, Jianhui He, Alian Liao, Deyu Han.

**Software**:Wanli Lu, Hongliang Zuo.

**Validation**: Linwei Yang, Jianhui He.

**Writing - original draft**: Linwei Yang, Xiaopeng Xu.

**Writing - review & editing**: Linwei Yang, Xiaopeng Xu.

## 6. Acknowledgements

This work was funded by the National Natural Science Foundation of China under grant No. 32273166, 32202974, 32573547 and 42376121; National Key Research and Development Program of China 2022YFF1000304; Natural Science Foundation of Fujian Province, China 2024J01432; Earmarked fund for China Agriculture Research System CARS48. Natural Science Foundation of Guangdong Province, China 2023A1515010864.

---

**Fig S1.**
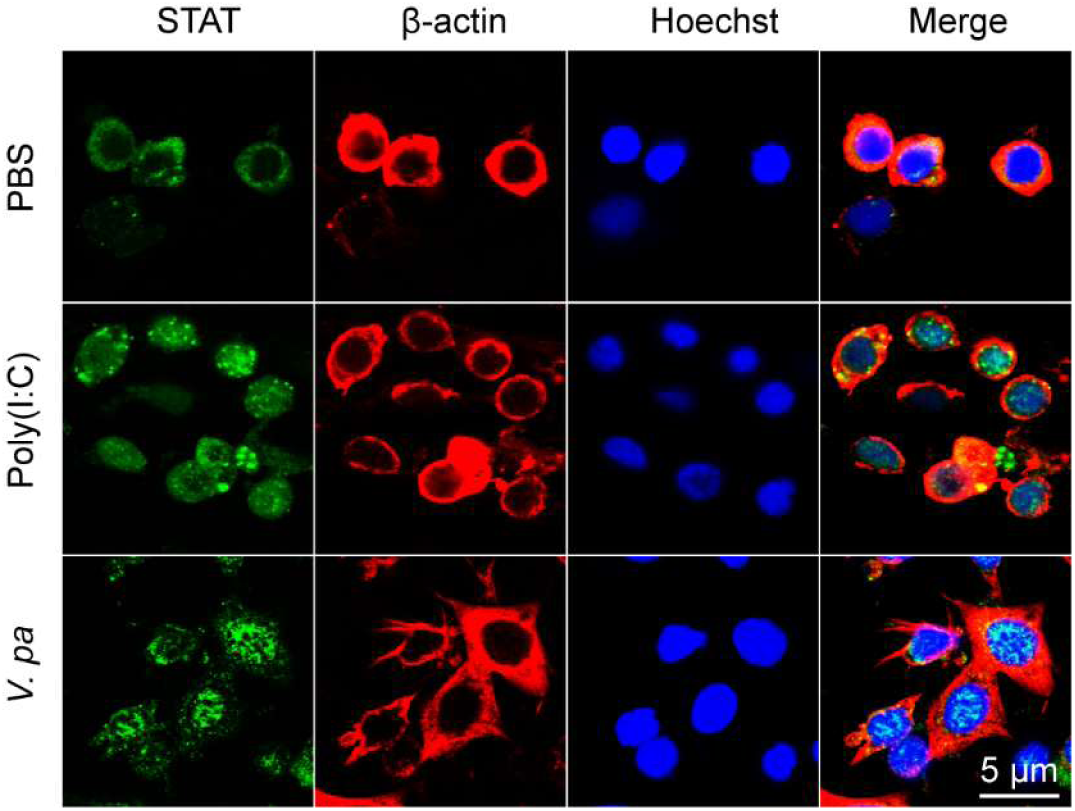
Immunofluorescence analysis of STAT nuclear translocation in shrimp hemocytes at 4 h post bacterial infection (5 × 10⁵ CFU) or Poly(I:C) stimulation (5 μg/shrimp). Cell nuclei were stained with Hoechst. Bar = 5 μm.

**Fig S2.**
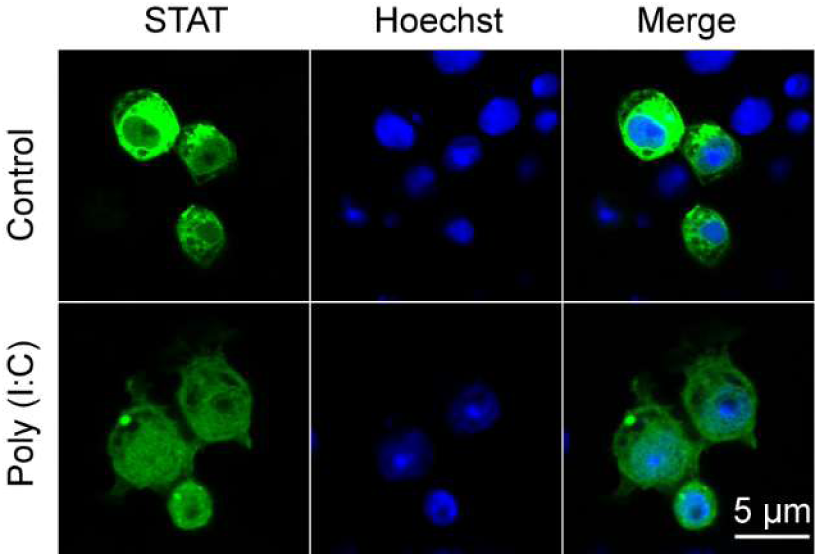
Immunofluorescence analysis of STAT nuclear translocation in S2 cells following stimulation with 0.5 μg/mL poly(I:C). Cell nuclei were stained with Hoechst. Bar = 5 μm.

**Fig S3.**
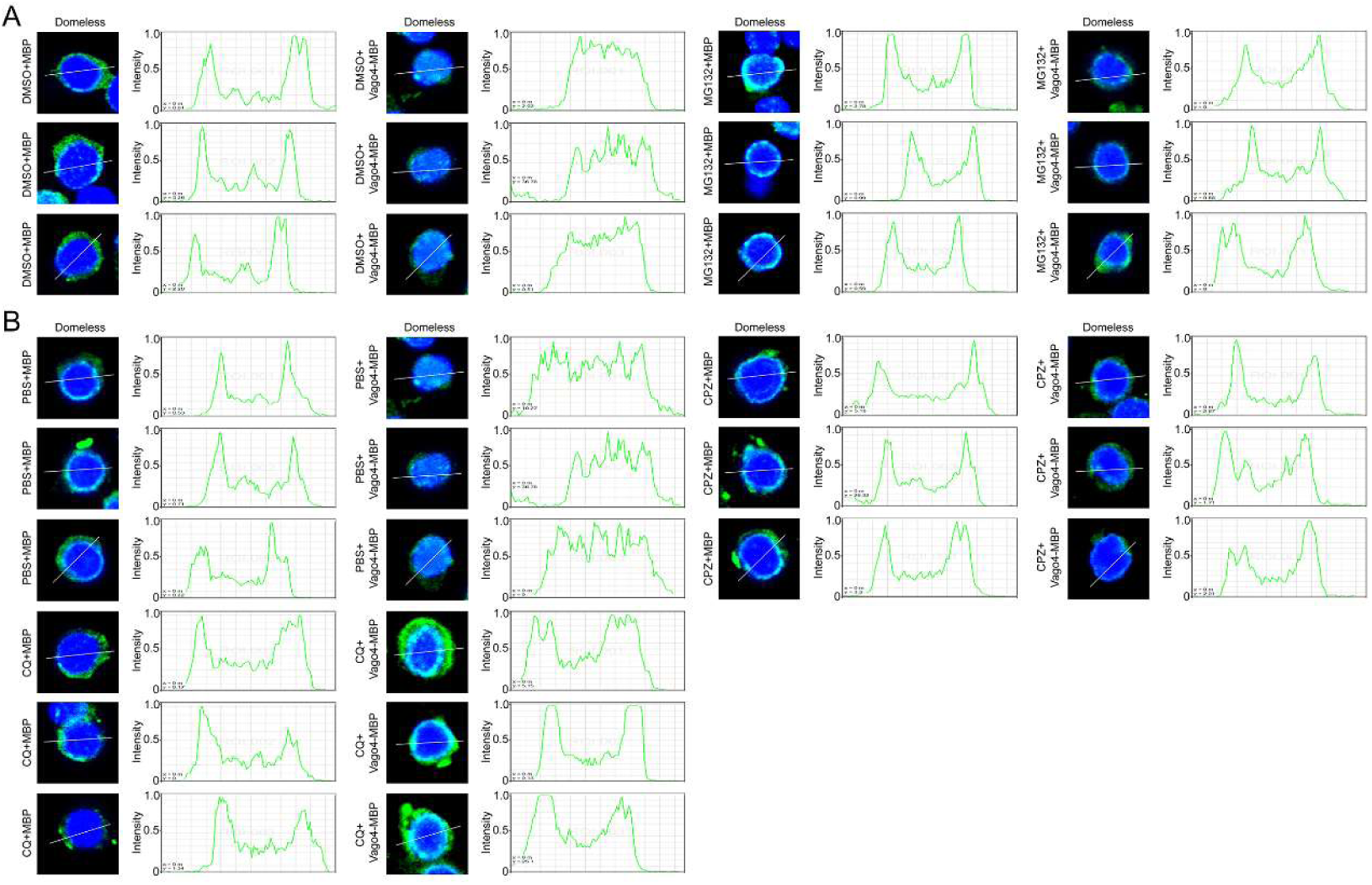
(A) Immunofluorescence analysis of Domeless protein levels in shrimp hemocytes 4 h after co-injection of recombinant Vago4 protein (10 μg) with either MG132 (20 μg), or DMSO. (B) Immunofluorescence analysis of Domeless protein levels in shrimp hemocytes 4 h after co-injection of recombinant Vago4 protein (10 μg) with either chloroquine (CQ, 100 μg), chlorpromazine (CPZ, 50 μg) or PBS. Redistribution of Domeless from the plasma membrane to the cytoplasm was analysis using SP8 Leica laser confocal microscopy imaging system (SP8, Leica, Germany). Cell nuclei were stained with Hoechst. Bar = 10 μm.

**Fig S4.**
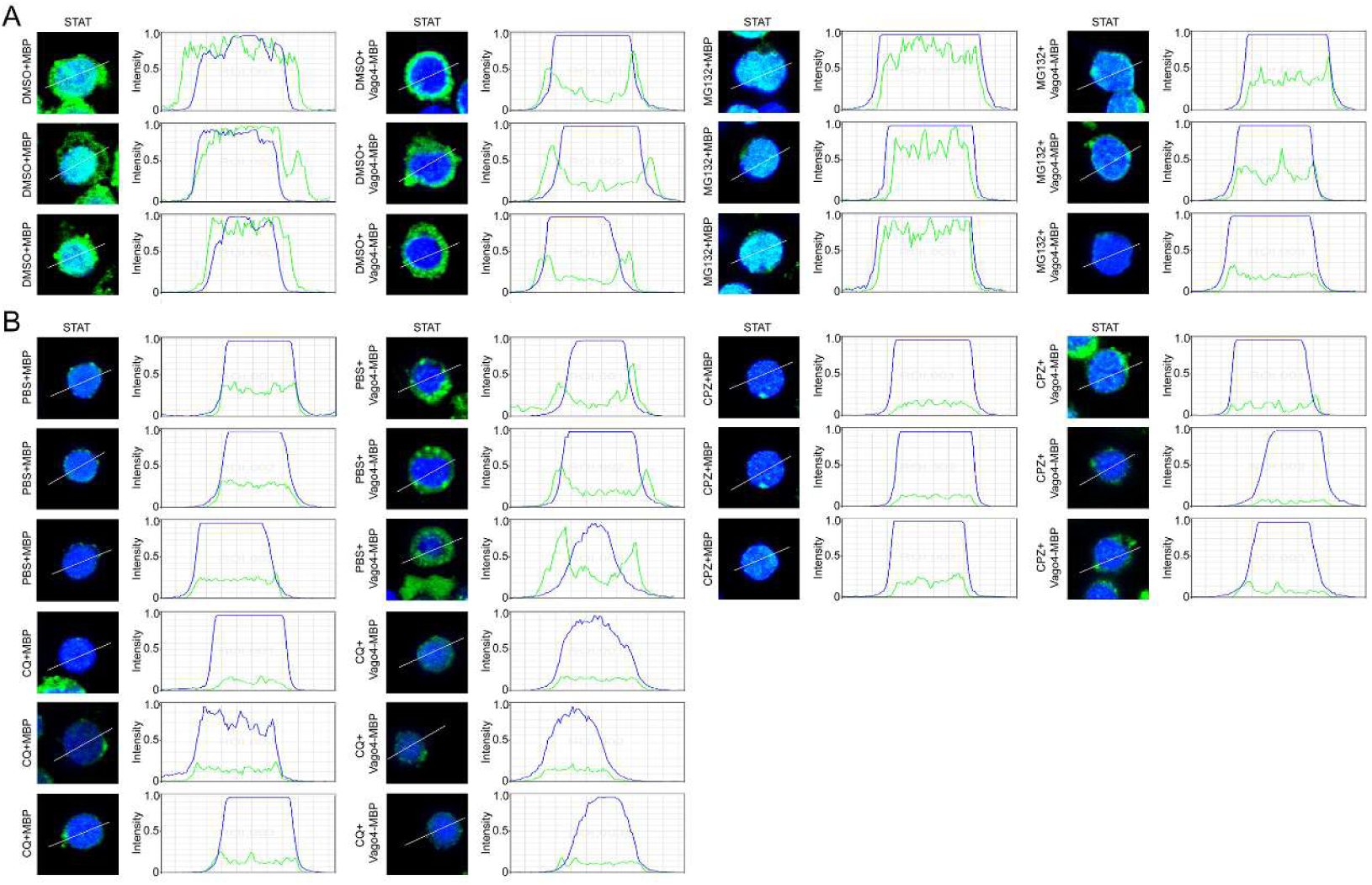
(A) Immunofluorescence analysis of STAT nuclear translocation in shrimp hemocytes 4 h after co-injection of recombinant Vago4 protein (10 μg) with MG132 (20 μg), or DMSO. (B) Immunofluorescence analysis of STAT nuclear translocation in shrimp hemocytes 4 h after co-injection of recombinant Vago4 protein (10 μg) with either chloroquine (CQ, 100 μg), chlorpromazine (CPZ, 50 μg) or PBS. Co-localization of STAT (green) and Cell nuclei (blue) were analysis using SP8 Leica laser confocal microscopy imaging system (SP8, Leica, Germany). Cell nuclei were stained with Hoechst. Bar = 10 μm.

## Notes

### Competing Interest Statement

The authors have declared no competing interest.

